# Comparative analysis of variation and selection in the HCV genome

**DOI:** 10.1101/078584

**Authors:** Juan Ángel Patiño-Galindo, Fernando González-Candelas

**Author notes:** Address for correspondence: Prof. Fernando González-Candelas, Unidad Mixta Infección y Salud Pública FISABIO-CSISP/Universitat de Valéncia – Instituto Cavanilles de Biodiversidad y Biología Evolutiva c/ Catedrático José Beltrán, 2, 46980 Paterna, Valencia. SPAIN.

## Abstract

Genotype 1 of the hepatitis C virus (HCV) is the most prevalent of the variants of this virus. Its two main subtypes, HCV-1a and HCV-1b, are associated to differences in epidemic features and risk groups, despite sharing similar features in most biological properties. We have analyzed the impact of positive selection on the evolution of these variants using complete genome coding regions, and compared the levels of genetic variability and the distribution of positively selected sites. We have also compared the distributions of positively selected and conserved sites considering different factors such as RNA secondary structure, the presence of different epitopes (antibody, CD4 and CD8), and secondary protein structure. Less than 10% of the genome was found to be under positive selection, and purifying selection was the main evolutionary force in both subtypes. We found differences in the number of positively selected sites between subtypes in several genes (*Core*, HVR2 in *E2, P7*, helicase in *NS3* and *NS4a).* Heterozygosity values in positively selected sites and the rate of non-synonymous substitutions were significantly higher in subtype HCV-1b. Logistic regression analyses revealed that similar selective forces act at the genome level in both subtypes: RNA secondary structure and CD4 T-cell epitopes are associated with conservation, while CD8 T-cell epitopes are associated with positive selection in both subtypes. These results indicate that similar selective constraints are acting along HCV-1a and HCV-1b genomes, despite some differences in the distribution of positively selected sites at independent genes.

## Introduction

Hepatitis C Virus (HCV) is the main causal agent of non-A non-B viral hepatitis, a pandemic with a global prevalence of 2.8%, affecting more than 185 million people worldwide (Mohd Hanafiah et al. 2013). Its high prevalence and the possibility that the virus causes chronic infections, which occur in about 80% of the cases, ending in hepatocellular carcinoma make HCV a major cause of concern for public health. HCV belongs to the genus *Hepacivirus* in the *Flaviviridae* family and its single, positive-sense, single-strand RNA genome of 9.6 kb encodes a polyprotein of more than 3000 amino acids (Takamizawa et al. 1991). This polyprotein is cleaved into 3 structural (Core, Envelope 1(E1), E2) and 7 non-structural (P7, NS2, NS3, NS4A, NS4b, NS5A and NS5b) proteins by means of both viral and human proteases (reviewed in Lindenbach and Rice 2005).

Like most RNA viruses, HCV is highly variable genetically and is phylogenetically divided into seven genotypes (named from 1 to 7) (Smith et al. 2014) which present more than 30% divergence at the nucleotide level among them. Most genotypes are further divided into subtypes, with 20-25% divergence among them (Simmonds et al. 1993). Genotype 1 is the most prevalent variant (Messina et al. 2015) and also the one with the lowest sustained viral response (SVR) rate to treatment with pegylated interferon and ribavirin (Pang et al. 2009), still the most frequent therapeutic option for hepatitis C infection. More than 10 HCV-1 subtypes have been reported so far (Kuiken et al. 2005). Among them, subtypes 1a and 1b are the most prevalent and cause about 40% of the total infections by the virus.

These two viral variants present some differences. There are epidemiological differences between their major transmission groups and these influence their phylodynamics. HCV-1a has been historically associated with transmission among intravenous drug users and HCV-1b with transfusions and other nosocomial and community transmissions (Shepard et al. 2005). Although subtype 1b apparently appeared around 10 years later than subtype 1a, it started an explosive growth phase 20 years earlier (in the 1940s), coincident with the start of widespread use of blood and blood derivatives in transfusions (Magiorkinis et al. 2009). Clinical differences between these two subtypes may also exist: sustained viral response (SVR) to treatment with pegylated interferon and ribavirin has been reported to be significantly higher in subtype 1a than in 1b (Pellicelli et al. 2012). Finally, at the genetic level, Torres-Puente et al. (2008) reported differences in genetic variability between these two subtypes, with HCV-1b showing higher nucleotide diversity in genes *E1, E2* and *NS5A.*

Most published studies detecting positively selected sites in this virus have analyzed only the *E1-E2* and/or *NS5A* genes (Cuevas et al. 2009; Cuevas et al. 2008; Humphreys et al. 2009; Sheridan et al. 2004). This is due to the relevance of the proteins that they encode in the viral response to the immune system and to treatment and, in consequence, for the establishment of persistent infection. Similar studies using complete coding regions published so far (Campo et al. 2008; Cannon et al. 2008) have used methods for the inference of positive selection that lack power, as they are based on estimating the ratio of nonsynonymous to synonymous substitution rates (dN/dS) from the reconstruction of ancestral states (Nei and Gojobori 1986). Hence, we currently lack a detailed view of the action of selection as a factor in the evolution of the HCV genome and how it may differentially affect major variants of this virus.

Our goal in this work is to analyze the impact of positive selection on the evolution of the genomes of these two viral variants, comparing the selective pressures in the different proteins of these two subtypes. For this, we present analyses with a comprehensive dataset of HCV-1a and 1b genomes and report a detailed comparative map of positively selected sites using, for the first time in HCV, the mixed effects model of evolution (MEME) (Murrell et al. 2012). This is a maximum-likelihood (ML) method that considers signals of both episodic (one that affects only to a subset of lineages) and pervasive (that affects to a large proportion of positively selected sites) selection more efficiently than other techniques. We have also estimated and compared the synonymous and non-synonymous substitution rates (dS and dN, respectively) and the heterozygosity (H) at individual sites. Finally, we have performed multivariate analyses in order to test the effects of RNA and protein structures and the presence of epitopes on the distribution of positively and conserved sites along their genomes.

## Results

In total, 415 sequences from HCV-1a and 204 sequences from HCV-1b were retrieved. After the application of additional criteria, the final data sets consisted of 393 HCV-1a and 179 HCV-1b sequences. The phylogenetic trees obtained with FastTree2 (GTR+GAMMA model) independently for each subtype, and that were used as input in the selection analyses, are shown in fig. 1.

**Figure 1.**
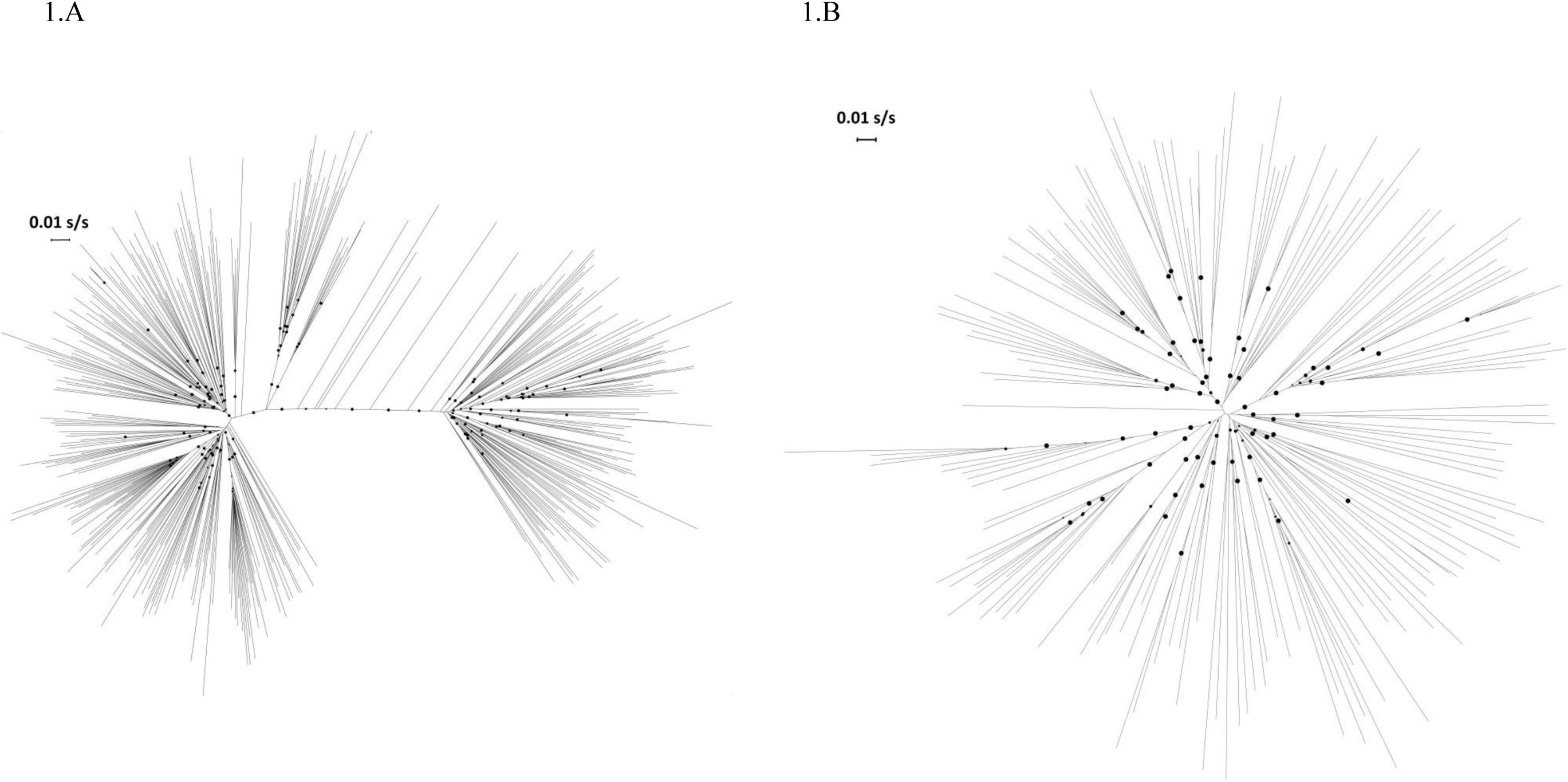
Phylogenetic trees of HCV-1a and HCV-1b, as obtained with FastTree2 (GTR+GAMMA model). Circles represent nodes with support ≥ 0.90.

FEL analyses detected 95 and 62 positively selected sites along the HCV-1a and HCV-1b genomes, respectively. In contrast, MEME analyses detected 315 (total dataset of HCV-1a) and 248 (HCV-1b) sites under positive selection. From those, 282 sites (HCV-1a) and 211 (HCV-1b) were not singletons (found only in one external branch) and were retained for ensuing analyses. All positively selected sites found by FEL were also found by MEME, and all these sites represented 9.4% and 7.0% of the total length of 1a and 1b protein encoding genomes, respectively. 99 sites were found to be positively selected in both subtypes. Additional details of these analyses are provided in Supplementary Table S1A and S1B (Supplementary Material online), including information on the genome location of each selected position, the frequency at which each selected site found in the full set of HCV-1a was also detected in the five subsets and LRT comparisons between MEME and FEL.

The performance of MEME and FEL for the detection of positively selected sites was statistically compared by performing LRTs of their log-likelihoods at each positively selected position detected by MEME, concluding that the mixed effects model outperforms the fixed effects model in most positions (209 in HCV-1a and 159 in HCV-1b), and could never be considered as a significantly worse method for the inference of positive selection in the datasets analyzed (Supplementary Tables S1A and S1B, Supplementary Material online).

Table 1 summarizes the distribution of positively selected sites along the different regions of the H77 reference polyprotein for the two HCV-1 subtypes and for the 5 random subsets of HCV-1a.

**Table 1.** Number positively selected sites, and its proportion (in brackets) at each gene of HCV-1a and HCV-1b. aa-number of amino acids encoded by each gene, n – number of sequences included in the dataset. Major domains are reported for E2 (HVR = Hyper variable region) and NS3 (PR = protease, HE = helicase) genes.

| Dataset | n | CORE<br>(191 aa) | E1<br>(192 aa) | E2<br>(363 aa) | P7<br>(63 aa) | NS2<br>(217 aa) | NS3<br>(631 aa) | NS4A<br>(54 aa) | NS4B<br>(261 aa) | NS5A<br>(448 aa) | NS5B<br>(591 aa) |
| --- | --- | --- | --- | --- | --- | --- | --- | --- | --- | --- | --- |
| HCV-1a | 393 | 10<br>(0.052) | 24<br>(0.125) | 70: 20 HVR1,<br>3 HVR2, 5 HVR3<br>(0.193) | 2<br>(0.032) | 28<br>(0.129) | 32: 9 PR,<br>23 HE<br>(0.051) | 0<br>(0.000) | 15<br>(0.057) | 50<br>(0.112) | 51<br>(0.086) |
| Subsets HCV-1a* | 197 | 5.8<br>(0.030) | 14<br>(0.073) | 52: 17.2 HVR1,<br>1.4 HVR2, 4.2 HVR3<br>(0.143) | 1.8<br>(0.029) | 17.2<br>(0.079) | 21: 7.2 PR,<br>13.8 HE<br>(0.033) | 0<br>(0.000) | 8.4<br>(0.032) | 36.8<br>(0.082) | 36.8<br>(0.062) |
| HCV-1b | 179 | 3 (h)<br>(0.015) | 16<br>(0.083) | 61: 18 HVR1, 6<br>HVR2, 6 HVR3<br>(0.168) | 5<br>(0.079) | 17<br>(0.078) | 27: 6 PR, 21<br>HE<br>(0.043) | 1<br>(0.019) | 8<br>(0.031) | 34<br>(0.076) | 39<br>(0.066) |

\*Results are given as the mean number of positively selected sites found in the five HCV-1a subsets.

A list of neutral sites was obtained from those sites that were not found to be under positive or negative selection in FEL or MEME and that were not totally conserved. We found that 231 in HCV-1a (7.7% of the total genome) and 271 positions in HCV-1b (9.0%) were evolving under neutrality (dN=dS). Of them, only 74 positions were coincident in both subtypes (Supplementary Table S1C, Supplementary Material online). Consequently, we estimated that there were 2465 and 2492 sites under purifying selection in HCV-1a and 1b, respectively.

Supplementary Figs. S1A and S1B (Supplementary Material online) show the phylogenetic trees obtained for HCV-1a and HCV-1b complete genome sequences, respectively. In these trees, the branches are colored according to the number of positively selected sites changing along each branch, as determined by parsimony using MacClade 4.08 (Maddison and Maddison 2005). Changes at positively selected sites were observed in both internal and external branches, although most changes accumulated in external branches.

Differences in the number of positively selected sites between proteins were found. The protein with the highest proportion of sites under positive selection was E2, followed by NS2 and E1. The proteins with the lowest proportion of sites under positive selection were NS4A and Core (Table 1). Whereas HCV-1b tended to present more positively selected sites in the second hypervariable region (HVR2) of E2 (mean = 1.4 sites in the subsets of HCV-1a, vs 6 sites in HCV-1b), in P7 (1.8 sites in HCV-1a vs 5 sites in HCV-1b) and the NS3-helicase (13.8 sites in HCV-1a vs 21 in HCV-1b). HCV-1a presented more positively selected sites only in Core (5.8 sites in HCV-1a, vs 3 in HCV-1b) (Table 1).

Mean and standard error (SE) estimates of dS, dN and heterozygosity (for both amino acids and nucleotides) obtained for the whole genomes, positively selected sites and neutral sites are available in Supplementary Tables S2A-C (Supplementary Material online), respectively.

After applying the FDR corrections, heterozygosity at third codon positions was found to be significantly higher in HCV-1b than in HCV-1a at the genomic level (HCV-1a: 0.161 ± 0.002 – mean ± standard error, HCV-1b = 0.181 ± 0.003; P = 1.3×l0^−8^). No significant differences were found in first or second codon positions nor amino acid sites (all P values > 0.05).

Mann-Whitney tests comparing heterozygosity between subtypes in all codon positions and amino acids and dS and dN of all positively selected sites revealed that HCV-1b presents significantly higher dN (HCV-1a: 1.257 ± 0.192, HCV-1b: 1.563 ± 0.271; P = 0.040) and heterozygosity at amino acids (HCV-1a: 0.239 ± 0.013, HCV-1b: 0.308 ± 0.016; P = 5.0×l0^−5^), first codon positions (HCV-1a: 0.155 ± 0.011, HCV-1b: 0.214 ± 0.013; P = 1.1×10^−5^), second codon positions (HCV-1a: 0.123 ± 0.010, HCV-1b: 0.169 ± 0.013; P = 0.003) and third codon positions (HCV-1a: 0.183 ± 0.010; HCV-1b: 0.206 ± 0.011; P = 0.023), but not significantly different dS (P = 0.90) (Supplementary Table S.2 B).

For neutral sites, no significant differences in dS, dN nor in heterozygosity between HCV-1a and HCV-1b were found, as concluded from the statistical tests (all P values > 0.05).

A map of the HCV-1a and HCV-1b genomes representing the different layers of data analyzed (conservation, positive selection, RNA structure, protein structure and epitopes for antibodies, CD8 and CD4 T cells) is shown in Fig. 2. In the logistic regression analyses, the models with the lowest AIC values were “RNA structure + CD4 epitope + CD8 epitope + Alpha helix” for HCV-1a and “RNA structure + CD4 epitope + CD8 epitope + Beta sheet” for HCV-1b. Comparing the distribution of positively and conserved sites at such layers of data revealed, for both HCV-1a and 1b, an association between conservation and secondary structure as well as with CD4 T cell epitopes. In contrast, an association between selection and CD8 T cell epitopes was found, although in this case the P-values for both subtypes increased to > 0.05 after the FDR correction (Supplementary Tables S3A and S3B, Supplementary Material online).

**Figure 2.**
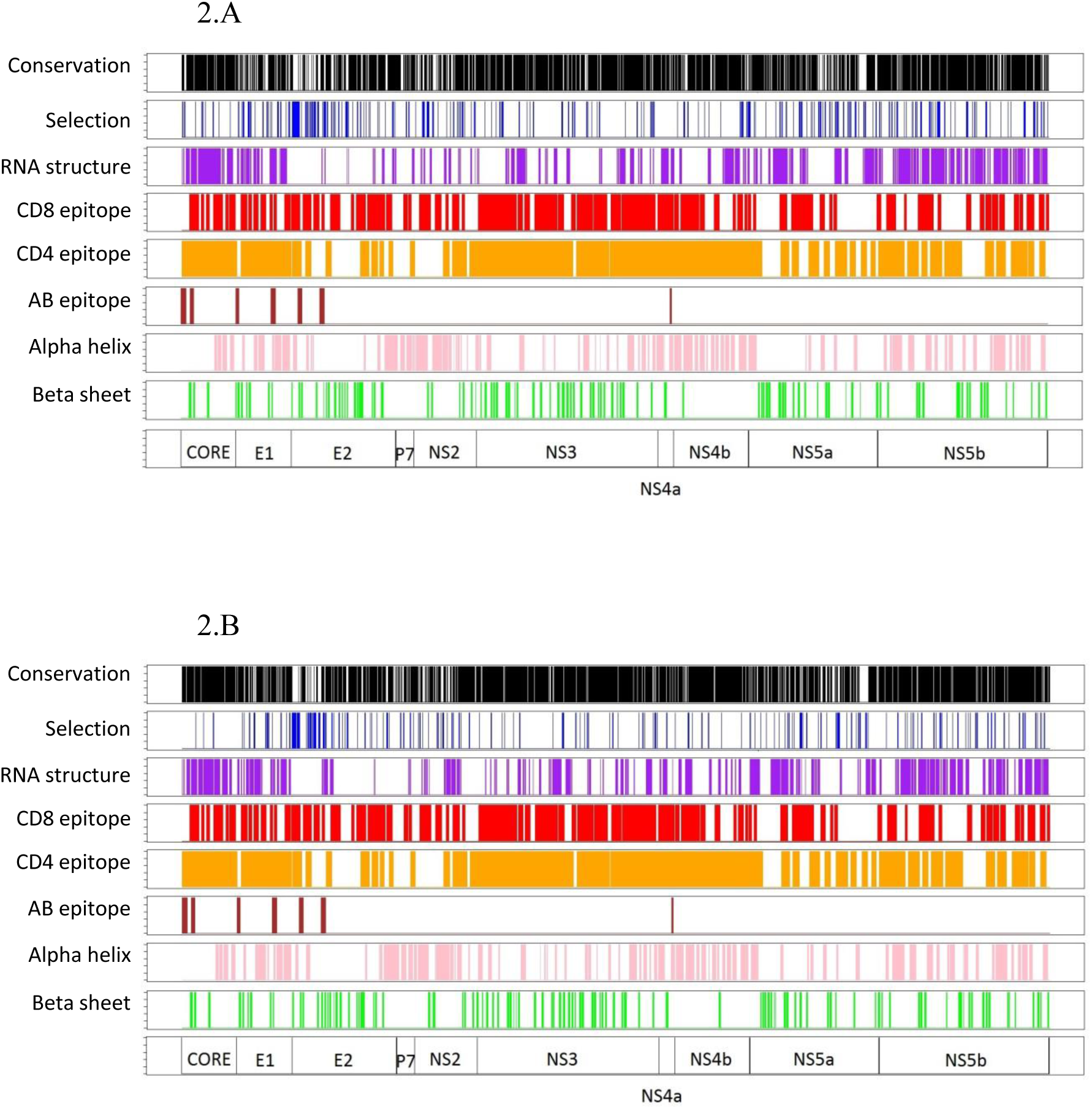
Map of the HCV-1a and 1b genomes, indicating the location of totally conserved amino acids (black), positively selected sites (blue) RNA secondary structures present in at least 50% of the sequences of each dataset (purple), CD8 T cell epitopes (red), CD4 T cell epitopes (orange), antibody(AB) epitopes (brown), alpha helixes (pink), beta sheets (green).

The same analyses were performed for all genes where at least 10 sites were found to be positively selected in both subtypes (E1, E2, NS2, NS3, NS5a and NS5b). In HCV-1a, an association between selection and the presence of CD8 epitopes was found NS2, although the P-value increased to > 0.05 after FDR correction. In subtype 1b, an association between conservation and presence of CD4 epitopes was found in NS3. Although associations between selection and CD8 epitopes, and between conservation and presence of beta sheets were found in E2 of HCV-1b, the P-value increased to > 0.05 after FDR correction (Supplementary table S.3A and S.3B).

## Discussion

In this work we have performed a comparative analysis of the evolutionary forces that shape the genomes of HCV-1a and HCV-1b, using both a fixed effects (FEL) and a mixed effects model (MEME) to detect sites evolving under different selective pressures: positive, purifying or neutral. Our results reveal several differences as well as similarities between these two relevant subtypes of HCV and help us understand the underlying factors driving their evolution at the genome level.

Previous studies analyzing selection in the HCV genome are more conservative than this one because they are based on Nei and Gojobori model (1986) for the calculation of dN/dS. Campo and colleges (2008) found a very similar number (n= 60) of sites evolving under positive selection along the HCV-1b genome using the SLAC method to those we found with FEL (n=62). However, our analysis with MEME detected more than 3 times more positively selected sites than these studies, in line with the expected increased power of this method (Murrell et al. 2012). In consequence, methods for the detection of positively selected sites that do not take into account that dN/dS can vary among lineages may underestimate the number of positively selected sites. This can occur when, for a given site, purifying selection prevails in most lineages, masking the detection of episodic positive selection occurring in a restricted number of lineages.

Most positively selected sites were mapped on the external branches of the phylogenetic trees for both viral subtype. This was expected, because the methods used to detect selection in fact identify sites under adaptive diversification when applied to the population or within-species level. Diversifying selection is known to be of major importance in the adaptive evolution of HCV, in which increasing global variability would be favored (Cuevas et al. 2009). Another possible explanation is that recent deleterious mutations, most of which are expected to be nonsynonymous, may not have been purged yet by selection and they will likely map on the external branches of the phylogeny. In consequence, for a given position different nonsynonymous substitutions might be detected as being positively selected when analyzing viral sequences from different patients when they actually represent transient polymorphisms. However, we excluded singletons from the list of positively selected positions, thus minimizing the presence of these false positives. The fact that there were no lineages accumulating a disproportionately large number of changes in the positively selected sites means that the results found in these analyses are representative of subtype, and not only of sub-clades.

We also found that purifying selection plays a major role in the evolution of HCV, with more than 2400 codons estimated to be under purifying selection in both subtypes. However, the number of neutral sites inferred from our work differs markedly from those detected by Campo et al. (2008) (833 sites). The difference might be explained by the different sampling sizes used in both studies (114 vs 179 HCV-1b sequences) but they are more likely due to differences in the methodology of both studies. The lower power of SLAC (Kosakovsky Pond and Frost 2005) used by Campo et al. (2008) to reject the null hypothesis of neutrality would be reflected in a large number of undetected, negatively selected sites. Furthermore, our criteria to detect neutral sites was more stringent, including only sites that were considered to be neither positively nor negatively selected in MEME and FEL and which presented H > 0 at the amino acid level, to avoid including codons under very strong, purifying selection for which no change could be detected.

We found positively selected sites in all HCV genes. Differences between subtypes in the distribution of positively selected sites were also detected. Whereas HCV-1b tended to present more positively selected sites in *E2* (HVR2), *P7* and *NS3* (helicase), HCV-1a presented more positively selected sites in the *Core* gene. This observation is based on the results obtained from the random subsets of HCV-1a, of similar number of sequences as that obtained for HCV-1b, in order to minimize the effect of sample size in the number of positively selected sites found. These subsets presented very similar levels of heterozygosity, dS, and dN when compared to the whole dataset of this subtype. Hence, differences between the two subtypes in these parameters are not likely due to their different sample sizes.

Differences in H between both subtypes were significant at the amino acid level and first, second and third codon positions of positively selected sites and only at third codon positions at the complete genome level. In contrast, differences in dN between the two subtypes were significant only for positively selected sites, whereas no significant differences in dS were detected in any of the comparisons performed. The different phylodynamic histories of HCV-1a and HCV-1b (Magiorkinis et al. 2009) could explain the higher genetic variability in third codon positions of HCV-1b at the genomic scale. HCV-1b started its explosive growth 20 years earlier than HCV-1a and its current effective infection size remains higher. This implies that it may have accumulated more genetic variability. Although no significant differences were found at first and second codon positions, where most changes are non-synonymous (a majority of these sites are totally conserved along the HCV genome), the significantly higher variability of HCV-1b with respect to HCV-1a was evident at third codon positions, in which most changes are synonymous. Interestingly, we did not find significantly higher dS in HCV-1b. According to the neutral theory of evolution, dS depends only on the neutral mutation rate but H is dependent also on the effective population size (Kimura 1983). Thus, we would expect HCV-1a and HCV-1b to present similar dS if their overall mutation rates are similar but to differ in H if they have different effective population sizes.

Despite the differences found in the distribution of positively selected sites, multivariate analyses at the genomic level revealed that HCV-1a and HCV-1b evolve under similar forces and constraints (RNA structure and CD4 T cell epitopes favoring conservation, and CD8 T cell epitopes favoring selection). Snoeck et al. (2011) mapped positively selected sites along the genome of HIV-1 and found that structured RNA and alpha helices in protein structure were associated with conservation whereas CD4 T-cell and antibody epitopes were associated with positive selection. However, they also found a significant association between CD8 and CD4 epitopes and conservation for several genes. Our results, regarding the overall association between secondary structure and conservation, are in line with those of Snoeck et al. (2011) and also with the results obtained by Mauger et al. (2015) with HCV. These authors suggested that conservation of secondary structures in this virus could facilitate persistent infection by masking the viral genome from its degradation by RNase L and innate antiviral defenses (Li and Lemon 2013; Washenberger et al. 2007).

In contrast to Snoeck et al. (2011) with HIV-1, in HCV we found no relevant association between antibody epitopes or protein structure and conservation or selection. Although the role of CD4 and CD8 T-cells in the immune response to HCV is well known, contrary to other viruses such as HIV or hepatitis B (Koziel 2005) there is not a clear pattern of antibodies response that distinguishes between recovery and chronic infection in HCV. Thus, further research to clarify their role in controlling HCV replication and to which extent such mechanisms determine the evolution of HCV is certainly needed.

Recently, Geller et al. (2016) estimated the *per site* mutation rate along the HCV genome, and found a small reduction in sites predicted to form base pairs. We performed a univariate analysis to check whether there was an unequal distribution between sites with low and high mutation rates, considering conserved and selected codons, and found no significant differences (Fisher’s exact test P > 0.20 in both subtypes).

The association between CD4 T-cell epitopes and conservation is remarkable. Given that epitopes are targets for the host immune system, it would be reasonable to expect epitopes to be under positive or diversifying selection (as for CD8), as an increased genetic variability would facilitate viral escape from the immune system. However, several studies have observed very conserved epitopes in HCV and other viruses (Lamonaca et al. 1999; Sanjuán et al. 2013; Sarobe et al. 2001; Snoeck et al. 2011). Sanjuán and colleagues (2013) suggested that HIV may take advantage of immune activation, thus favoring epitope conservation. Hence, if HCV also benefits from immune activation, the design of vaccines based on conserved epitopes would be detrimental.

After this manuscript was written, a paper by Cuypers et al (2016) was published, on September 6^th^, which also analyzed the different distributions of positively selected and conserved sites considering variables such as protein and RNA secondary structure and B-cell, CD8 and CD4 epitopes. We were unaware of this paper during the development of our work. They obtained similar results regarding the association between conservation and RNA structure and CD4 epitopes. However, some discrepancies were also obtained: they found an association between positive selection and CD8 epitopes only in HCV-1a, while we did not find intersubtype differences. In addition, they obtained significant associations between different protein structures (alpha helix and B-sheets) and conservation, whilst we did not find any significant association for these two variables. Although Cuypers et al (2016) used protein structure information derived from crystallized or nuclear magnetic resonance (NMS) when available, the extensive lack of such information along the HCV genome led both studies to predict such structures by means of computational methods. Consequently, differences in the methodology used for mapping protein structures, including the use of different protein structure prediction programs, may have caused these incongruent results.

In conclusion, we have produced a detailed map of positive selection along HCV-1a and 1b genomes and analyzed which variables can impose constraints or be associated to selection. We have shown that, despite purifying selection being the major evolutionary force acting on HCV, positive selection affects all genes along the HCV genome. Although there are differences in variability and the distribution of positively selected sites, both viral subtypes share similar selective pressures along their genomes. The results obtained from this study give information about the effect of some of the interactions between HCV and its host on HCV variability, which may be useful for antiviral research.

## Materials and Methods

Full coding regions from HCV-1a and HCV-1b genomes were retrieved form the VIPRBRC dataset on May, 2013 (Pickett et al. 2012) (amino acid positions 1 to 3011, according to the reference genome H77-GenBank accession number AF011753). Only sequences derived from human hosts were included. In addition, we ensured that all HCV sequences derived from DAA-naïve patients, excluding all sequences from DAA-treated patients described in the literature up to January 2015. Additional inclusion criteria were:

1. We included only one sequence per patient. For this, we performed a CD-Hit clustering analysis (Huang et al. 2010) with a similarity threshold of 0.95. Further information from the clustered sequences was obtained from the NCBI (http://www.ncbi.nlm.nih.gov/) in order to confirm that they were duplicate sequences.
2. Viral subtypes were confirmed using the COMET HIV-1/2 & HCV subtyping tool (Alcantara et al. 2009; De Oliveira et al. 2005, accessible at http://comet.retrovirology.lu).
3. Recombinant sequences were detected and subsequently removed using five different methods implemented in the RDP3 software: RDP, Geneconv, Bootscan, Maxchi and Chimera (Martin et al.,2005; Martin and Rybicki 2000; Martin et al. 2010; Padidam et al. 1999; Posada and Crandall 2001). The criterion to remove putative recombinant sequences was to obtain significant results with at least two different methods.

Multiple alignments were obtained for each HCV subtype independently using Muscle (Edgar 2004), as implemented in MEGA5 (Tamura et al. 2011).

Methods for the detection of positive selection usually become more powerful when the size of the dataset increases (unpublished results). To deal with the differences in size between the final datasets of HCV-1a and HCV-1b (393 and 179 sequences, respectively) we obtained 5 random subsets, each including half the total size of HCV-1a dataset (n=197), and performed the same positive selection analyses.

For each dataset, a phylogenetic tree was obtained with FastTree2 (Price et al. 2009), using a GTR+GAMMA evolutionary model, as recommended by the AICM analyses implemented in jModeltest (Posada 2008).

For each dataset and its corresponding tree, two different positive selection analyses were performed: (1) two-rate FEL (Fixed Effects likelihood), a maximum-likelihood (ML) method used to find independent sites under positive, neutral or purifying selection which considers that both dN and dS can vary between sites, while dN/dS remains constant along the different lineages of a given phylogenetic tree. It performs a likelihood ratio test (LRT) for each independent site to compare the null hypothesis of neutrality (dN = dS) with the alternative hypothesis of presence of either negative or positive selection (dN ≠ dS) (Kosakovsky Pond and Frost 2005); (2) MEME (Mixed Effects Model of Evolution), an extended version of FEL which considers that dN/dS can change across lineages. Unlike FEL, it models a variable dN/dS for each site across lineages using a two-bin random distribution with two categories of dN: Beta(-), for which dN ≤ dS and which occurs at a proportion q(-)of lineages, and Beta (+), in which dN is unrestricted when compared to dS (dN can be >, = or < than dS), which occurs at a proportion q(+) = 1 – q(−) of lineages. For each codon, it performs a LRT to compare the alternative hypothesis of the unrestricted Beta category with the nested null hypothesis of Beta (+) ≤ 1. This method has been reported to be more efficient in finding sites under positive selection than FEL and it has been recommended for finding both episodic and pervasive selection, with a type I error probability not higher than 0.05 (Murrell et al. 2012). Both analyses were performed with Hyphy (Kosakovsky Pond and Muse 2005) using the GTR model of nucleotide substitution and setting the significance level at 5%. Given that MEME and FEL are nested models, we compared their performance by calculating the LRT of their log-likelihoods at each codon (Murrell et al. 2012). According to a Chi-square distribution with 2 degrees of freedom, LRT values larger than 5.99 were considered to be significant.

Sites identified as being positively selected were mapped on the phylogeny and singletons were not considered for further analyses. This decision was taken in order to minimize the presence of false positives that could actually be sites in which a deleterious mutation had occurred but had not been removed from the viral population at the time of sampling. The remaining positively selected sites were mapped according to the H77 reference genome.

For each subtype, a list of neutrally evolving sites was also obtained by means of the following procedure:

1. For each HCV subtype, a list of sites not evolving under positive nor negative selection was obtained from the results of FEL, as neutral evolution is considered the null hypothesis.
2. Potentially neutral sites that were actually positively selected, as detected by MEME, were excluded.
3. Completely conserved sites were also excluded, as they were considered to be under very strong purifying selection. When a site has no variation, then the LRT from FEL or MEME does not have power to reject the null hypothesis of neutrality, but this site should not be considered as neutral.

Gene diversity (H) at each amino acid and nucleotide site (first, second and third codon positions) was calculated using the expression H=1−∑P_i_^2^ (where P_i_ is the frequency of each allele at a given site). dN, dS and H values were compared between subtypes using independent Mann-Whitney tests for: a) all genomic codons; b) positively selected codons and c) neutrally evolving codons. P-values obtained from non-independent statistical tests were corrected by means of False Discovery Rate corrections (FDR; (Benjamini and Hochberg 1995).

Frequencies of base-pairing at each nucleotide position for each HCV subtype were estimated with the software STRUCTURE_DIST (Tuplin et al. 2004), which analyses multiple RNA-folding patterns predicted by MFOLD (Zuker 2003). Antibody, CD8 and CD4 T-cell epitope positions were retrieved from the Los Alamos National Laboratory website (http://hcv.lanl.gov/content/immuno/immuno-main.html) and the Immune Epitope Database and Analysis resource (http://www.iedb.org) on June, 2015. Only human epitopes from HCV genotype 1 were included. Protein structures in both subtypes were inferred with JPred4 (Drozdetskiy et al. 2015), which predicts the location of secondary structures of proteins (alpha helixes, beta sheets) from multiple alignments of protein sequences. All these sites were mapped according to the H77 reference genome.

Logistic regression analyses (general linear model, GLM) were performed to compare, in each subtype, the distribution of positively selected and conserved positions. Several binary variables at each position were considered in the linear models: (1) RNA base-pairing (consensus ≥ 0.50) at the subtype level, (2) CD8 T-cell epitope, (3) CD4 T-cell epitope, (4) antibody epitope, (5) alpha helix, and (6) beta sheet. Positions were considered to be conserved if they had H = 0 at the amino acid level. An initial model, which included all the variables, was built. Stepwise model selection by Akaike Information Criterion (AIC) was performed with the R package “MASS” (Ripley et al. 2012) with the aim of including only relevant predictors. Stepwise search was performed in both directions (which tests at each step for variables to be included or excluded), and the best quality model for the GLM was chosen as the one with the lowest AIC value. P-values obtained in the logistic regression analyses were corrected by means of FDR. All statistical tests were performed as implemented in R (R Core Team 2015).

## Acknowledgements

This work was supported by projects BFU2011-24112 and BFU2014-58565-R from Ministerio de Economía y Competitividad (Spain). JAPG is recipient of a FPU fellowship from Ministerio de Educación y Ciencia (Spain).

## Supplementary material

**Supplementary Tables S1.** Positively selected and neutrally evolving sites in HCV-1a and 1b with comparison between the results obtained with MEME and FEL methods.

**Supplementary Tables S2.** Estimates of H, dN and dS for total genome, positively selected and neutrally evolving sites in HCV-1a and 1b for the different protein coding genes in the HCV genome.

**Supplementary Table S3.** Summary of logistic regression analyses for factors associated to positive selection in protein coding genes of HCV-1a and 1b subtypes.

**Supplementary Fig. S1.** Trees obtained with FastTree2 (model GTR + GAMMA), in which branch lengths and colors depend on the number of unambiguous changes in positively selected sites. Fig. S1A: HCV-1a. Fig. S1B: HCV-1b.

